# The Interaction of Context Constraints and Predictive Validity during Sentence Reading

**DOI:** 10.1101/2022.09.21.508808

**Authors:** René Terporten, Eleanor Huizeling, Karin Heidlmayr, Peter Hagoort, Anne Kösem

**Affiliations:** Donders Centre for Cognitive Neuroimaging; Max Planck Institute for Psycholinguistics; UMR 7114 MoDyCo, CNRS-Universite Paris Nanterre; Centre de Recherche en Neurosciences de Lyon

## Abstract

Words are not processed in isolation, instead they are commonly embedded in phrases and sentences. The sentential context influences the perception and processing of a word. However, how this is achieved by brain processes and whether predictive mechanisms underlie this process remains a debated topic. To this end we employed an experimental paradigm in which we orthogonalized sentence context constraints and predictive validity, which was defined as the ratio of congruent to incongruent sentence endings within the experiment. While recording electroencephalography, participants read sentences with three levels of sentential context constraints (high, medium and low). Participants were also separated into two groups, which differed in their ratio of valid congruent to incongruent target words that could be predicted from the sentential context. For both groups we investigated modulations of alpha power before, and N400 amplitude modulations after target word onset. The results reveal that the N400 amplitude gradually decreases with higher context constraints. Contrary, alpha power is non-monotonically influenced, displaying the strongest decrease for high context constraints over frontal electrode sites, while alpha power between medium and low context constraints does not differ. This indicates that both neural correlates are influenced by the degree of context constraint but are not affected by changes in predictive validity. The results therefore suggest that both N400 and alpha power are not unequivocally linked to the predictability of a target word based on larger contextual information.

## INTRODUCTION

In daily language usage, words are not processed in isolation, but they are commonly embedded in phrases or sentences. It is known that sentences create a context which even biases the perception and processing of a word. For instance, contextual information processed during sentence reading is known to facilitate the processing of new linguistic input (Kuperberg & Jaeger, 2016). While this phenomenon is well documented (DeLong et al., 2005; Frank et al., 2015; Freunberger & Roehm, 2017; Ito et al., 2016; Kutas & Federmeier, 2011; Van Petten & Luka, 2012), the mechanisms at the neurobiological origins of the processing of sentential linguistic information are still debated (Huettig, 2015; Huettig & Guerra, 2019; Nieuwland et al., 2019). On the one hand, the effects of context constraints could occur incrementally via integration mechanisms, which consist in integrating the (bottom-up) activated word meaning with its context *upon* its presentation (Bar, 2007; Gerrig & McKoon, 1998; Huettig, 2015; Lau et al., 2012). Conversely, the processing of contextual information could result from neurobiological mechanisms that support linguistic prediction (Federmeier et al., 2007): based on the contextual information, the brain would build predictions about certain linguistic features of the incoming words *prior* to the arrival of the sensory evidence.

Whether and to what degree the brain employs predictive mechanisms, during language processing that is influenced by sentential constraints, remains an outstanding question. Linguistic prediction has received experimental support. However, one way to answer the core question is to experimentally dissociate sentential context and linguistic predictive validity, which refers to the extent to which contextual information can be used to engage in linguistic predictive processes. In line with this idea, a recent discussion (Huettig, 2015) pointed to the possibility that distinct contextual processes could potentially be recruited depending on the task specific goal of the participant. Evidence from semantic priming tasks indicate that the amplitude of the N400 evoked response to a target word is not only dependent on the prime word but also on the experimental setup, in this case the proportion of valid primes in the design (Brown et al., 2000; Holcomb, 1988; Lau et al., 2012). Specifically, Lau and colleagues (2012) demonstrated that changes in predictive validity modulated the N400 amplitude for highly predictable word pairs. Yet, semantic priming paradigms rely on associative mechanisms (Megan A. Boudewyn et al., 2012; Brothers et al., 2017; Kuperberg et al., 2010; Lau et al., 2012), whereas the instantiation of sentential context requires the combinatorial operation of individual word meanings in a sentence.

Going beyond word pair processing, different explicit task instructions for sentence reading influence the effect of context constraint onto brain responses: the N400 amplitude at the target word was stronger when participants were explicitly asked to predict the word, compared to when they were asked to understand the sentence (Brothers et al., 2017, 2019). Here the cloze probability of the target words and their predictive validity were manipulated, while sentence context constraints were held constant. However, a successful attempt to dissociate sentential context and predictive validity from each other would require a design in which both factors - sentence context constraint and predictive validity - vary orthogonally.

Another approach to find evidence for prediction in language processing is to pay attention to a period *before* the processing of a target word. A neural marker that is sensitive to variations in sentential context before target word occurrence could likely be linked to underlying predictive processes. Brain oscillatory responses in the alpha (8-12 Hz) frequency range have been linked to linguistic predictive mechanisms prior to the occurrence of a target word (Lam et al., 2016; Piai et al., 2017; Rommers et al., 2017; Wang et al., 2017), with decreases in alpha power relating to the processing of sentence context constraints, which in turn contribute to language prediction. The power decrease has been found to be stronger for sentential context constraints that lead to a strong prediction of a target word as compared to when its predictability is very low (M. Bastiaansen & Hagoort, 2015; Piai et al., 2017; Rommers et al., 2017; Wang et al., 2017; Willems et al., 2008), though the direct link between alpha power and linguistic predictability has been challenged in a recent report (Terporten et al., 2019). Terporten and colleagues (2019) used varying degrees of sentential constraints to influence linguistic predictability. Against initial expectations, alpha power before target word occurrence has been found to be non-monotonically related to the strength of sentential context constraints. While being able to identify alpha oscillations as suitable neural marker to identify language prediction processes, predictive validity was not the target of the manipulation.

The current study attempts to replicate the findings form Terporten et al. (2019). Furthermore, it is tested whether alpha oscillations are related to linguistic prediction by investigating whether they are sensitive to the predictive validity of a sentential context. Predictive validity is defined here as the ratio of congruent to incongruent sentence endings within the set of sentences presented to the participants. For a set of sentences, the influence of linguistic predictive validity is independently manipulated from varying degrees of sentential context constraints. Participants passively read sentences with either a high (HC), medium (MC) or low (LC) context constraints. The validity of these predictions was manipulated experimentally using different environments. Specifically, participants were split into two groups: one group receiving mainly (80%) congruent target words to the previously established sentence context, the other group receiving mainly (80%) incongruent target words. The proportion of valid predictions was thought to alter the way of how sentence context constraints were evaluated to generate linguistic predictions (Bosker et al., 2019; Lau et al., 2012). We observed the influence of sentence context constraint and language predictive validity on two distinct neural markers, the N400 and alpha (8 Hz-12 Hz) oscillations, recorded with electroencephalography (EEG). We investigated how the N400 amplitude at target word onset, and pre-target word alpha power were modulated by context constraints and by the predictive validity of the target words. We expected the N400 amplitude to be gradually influenced by sentence context constraints, with higher constraints resulting in a reduction in amplitude (less negative). We also explored later post-N400-positivities (PNP) as an event related effect that follows the N400, which has been shown to be sensitive to semantic plausibility and predictability (Van Petten & Luka, 2012). Based on our earlier approach (Terporten et al., 2019) we expected pre-target alpha power to be modulated by sentence context constraints, but we expected alpha desynchronization to be non-monotonically linked to the predictability of the target word. If a non-monotonic modulation of alpha power before target word onset is also mediated by predictive validity, the results would be in favor of the brain employing predictive mechanisms during the encoding of sentential context.

## MATERIALS & METHODS

### Participants

In total, 70 participants were invited from the participant pool of the Max Planck Institute (MPI) for Psycholinguistics, Nijmegen. All participants gave their informed written consent in accordance with the declaration of Helsinki, and the local ethics committee (CMO region Arnhem-Nijmegen). All participants were Dutch native speakers, right-handed, had normal or corrected-to-normal vision and did not suffer from neurological impairment or dyslexia. After completion of the experiment, the participants received 18 euro. One participant did not finish the experiment and was excluded such that 69 participants (mean age 25 years, range 19-41; 20 males) were included for the analyses.

### Stimulus material

The stimulus set used in this study consisted of 203 critical sentence triplets from Terporten et al. (Terporten et al., 2019). While the original stimulus set contained only congruent sentence endings, an additional set of sentences was created with only incongruent sentence endings (see table 1 for examples). This approach resulted in two sets of sentence triplets: a congruent and an incongruent set of sentence triplets. Manipulations in predictive validity were achieved by changing the ratio of congruent to incongruent sentence endings for the final experimental set of sentence triplets. Each sentence for both groups belonged to either a high context (HC), medium context (MC), or low context (LC) constraining condition. The degree of constraint for a given sentence was manipulated by changing one word, the *context constraining word*. This word was always at the same position within a sentence with regard to a triplet (table 1). Across the conditions, these context constraining words were matched with regard to word length (F(2, 606) = 0.78, p = .457, with a Mean (SE) of HC: 7.12 (2.26); MC: 7.1 (2.54); LC: 7.37 (2.61)) and word frequency (F(2, 584) = 1.98, p = .138, with Mean (SE) of HC: 2.4 (0.78); MC: 2.56 (0.87); LC: 2.5 (0.84); based on the Dutch SUBTLEX-NL database (Keuleers et al., 2010). The degree of context constraints was measured in Terporten et al. (2019) by using a sentence completion task in which participants had to fill in a missing target word based on a preceding sentence context. Sentences with the same target words filled in by the participants received higher context constraint ratings. The degree of context constraints differed significantly between constraining conditions (F(2, 606) = 442.84, p < .001). HC sentences showed the strongest degree of context constraints (Mean (SE) = 77% (17.74)), followed by MC (Mean (SE) = 50% (18.67)) and LC (Mean (SE) = 28% (11.97)). In (Terporten et al., 2019) all sentence final words - the *target words* – were possible continuations of the preceding context (congruent target words). The cloze probabilities of the congruent target words differed significantly between conditions (F(2, 606) = 468.16, p < .001), with HC showing the highest cloze probability (Mean (SE) = 77% (17.74)), followed by MC (Mean (SE) = 42% (25.94)) and LC (Mean (SE) = 15% (15.82)). Measures of context constraints highly correlated with measures of cloze probability for *congruent* target words (r = 0.93, *p* < .001). In addition to the congruent stimulus set, a stimulus set was created with 203 *incongruent* target words. The congruent and the incongruent stimulus set differed significantly from each other on pre-tested ratings of plausibility (F(1,1312) = 4772.23, *p* < .001). The incongruent target words did not occur in the pretest of the congruent stimulus set and therefore all have a cloze probability of 0%. Congruent and incongruent target words were matched on word length (t(404) = −1.12, *p* = 0.264, with a Mean (SE) of Congruent: 5.79 (0.14); Incongruent: 6.0 (0.13)) and word frequency (t(404) = 1.29, *p* = .199, with a Mean (SE) of Congruent: 3.07 (0.05); Incongruent: 2.98 (0.04)); based on the Dutch SUBTLEX-NL database (Keuleers et al., 2010).

**Table 1:**
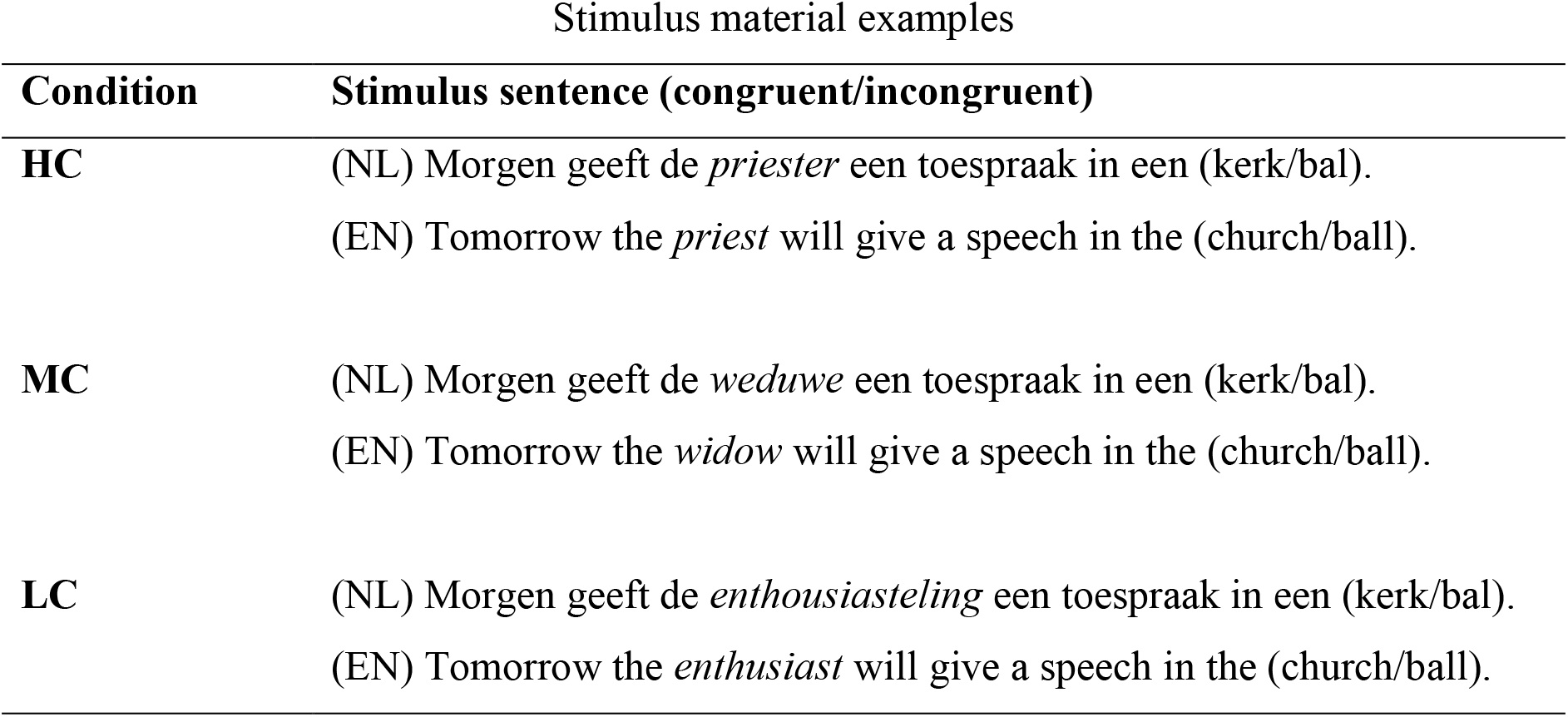
Example Dutch sentence triplet from the final stimulus set with its English translation. The context constraining conditions – high (HC), medium (MC) and low (LC) constraints - were manipulated by changing one context constraining word.

| Condition | Stimulus sentence (congruent/incongruent) |
| --- | --- |
| HC | (NL) Morgen geeft de *priester* een toespraak in een (kerk/bal).<br>(EN) Tomorrow the *priest* will give a speech in the (church/ball). |
| MC | (NL) Morgen geeft de *weduwe* een toespraak in een (kerk/bal).<br>(EN) Tomorrow the *widow* will give a speech in the (church/ball). |
| LC | (NL) Morgen geeft de *enthousiasteling* een toespraak in een (kerk/bal).<br>(EN) Tomorrow the *enthusiast* will give a speech in the (church/ball). |

A practice stimulus set was also created, including a selection of 50 sentence in total, split in congruent and incongruent sentences from (Wang et al., 2017). Half of the sentences were defined as HC, while the other half was defined as LC for each congruency condition separately (see (Wang et al., 2017) for details). For the EEG experiment, six counterbalanced lists were created. Three of these lists contained 80% of congruent target words, while the other three lists contained 80% of incongruent target words. The practice stimulus set was thought to bias participants’ expectation of the predictive validity of the context constraints, towards the respective proportion of (in)congruent target words in the critical stimulus set. For all lists, the three levels of context constraints were pseudo-randomly distributed across the set.

### Experimental procedure

Participants were comfortably seated in front of a screen in a dimly illuminated room. They were instructed to rest their right arm on the table in front of them and to access a button box with their right hand. At 70 cm and with a 25°-35° viewing angle, a screen was located to which all stimuli were projected. Written stimuli were shown in black, on a grey background.

Participants were instructed to silently read a word-by-word display of sentences on the screen, and to focus on the content of each sentence. It was explained that sometimes (after 25%of the sentences; subjects were not informed about the precise percentage) a question would be prompted about the content of the previously displayed sentence. The participants were required to answer this question with ‘yes’ or ‘no’ by button press. The answer possibilities (‘yes’/’no’) were displayed randomly on the left or right side of the screen and a left or right button had to be pressed accordingly. The occurrence of these questions throughout the experiment was at random intervals. The goal of these questions was to ensure that the participant kept processing the content of the sentence material throughout the experimental session.

A trial began with the presentation of a fixation cross in the middle of the screen for 500 ms. This was followed by a blank screen for a jittered interval of 500-1200 ms. The sentences were presented word-by-word. Each word was displayed for 200 ms, followed by a blank screen of 800 ms. The inter stimulus interval of 1000 ms was chosen to avoid the influence of the evoked response from stimulus onset onto pre-target alpha activity. Another blank screen occurred for 2000 ms (Fig. 1) after (sentence final) target word offset. In 25% of the cases, a catch question was displayed, with the full question centered on the screen and the yes-no answers randomly split to the left or right side.

**Figure 1:**
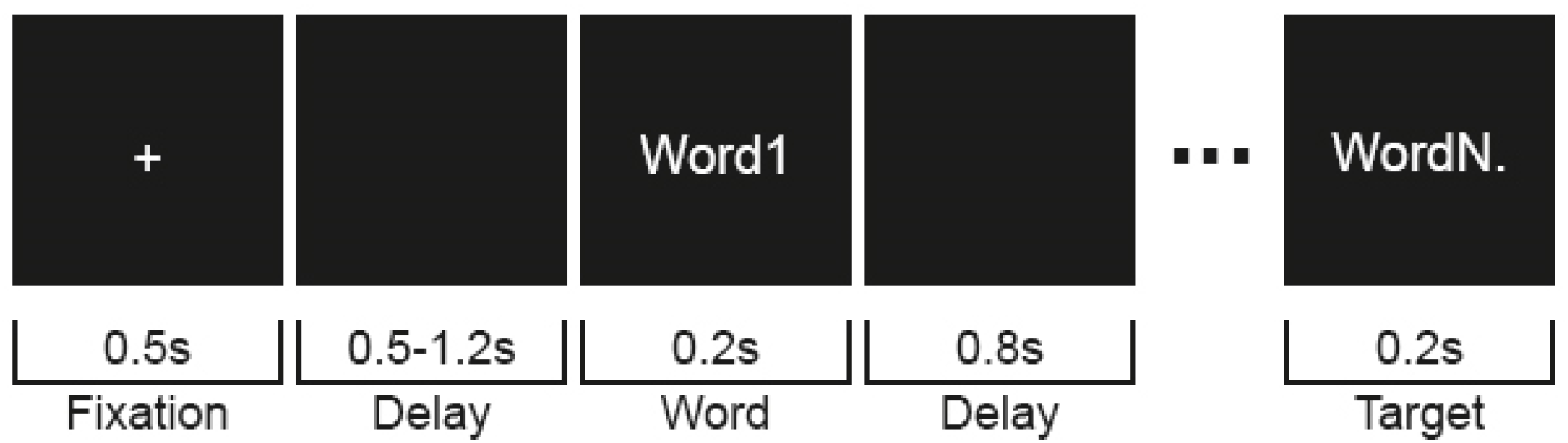
A schematic display of a trial procedure showing the duration of each screen. A trial began with the display of a fixation period, followed by a blank screen. Subsequently the sentence was visually displayed by a word by word presentation, up to the final word as indexed by the period. Between words, a black screen served as delay before a subsequent word was shown.

Each participant was presented with either a congruent or incongruent practice set, followed by the corresponding (low- or high predictive-validity) experimental list. The order of experimental lists across participants was pseudo-randomized. This resulted in half of the participant pool belonging to a high predictive validity group, while the other half belonged to a low predictive validity group. For each group, participants were presented with 50 practice sentences from Wang et al. (Wang et al., 2017) at the start of the experiment to prime the statistics of the experimental predictive validity. This was followed by 203 critical sentences (203 trials) which were presented in a random order. Trial presentation was divided into four blocks, separated by self-paced breaks in-between. In total, the experimental procedure took 60 min.

### Data acquisition

The participants’ electroencephalogram (EEG) was recorded online. A custom actiCAP 64-electrode montage (Brain Products, Munich, Germany) was used, with 58 equidistant electrodes mounted in the cap. Four electrodes measured EOG, with two horizontal EOG electrodes placed next to the left and right eye. Vertical EOG was measured by placing two electrodes above and below the participant’s left eye. The reference electrode was placed on the left mastoid. The ground electrode was placed on the forehead, above the nasion. Data were filtered online with a high pass filter of 0.02 Hz and a low pass filter of 500 Hz.

### Data preprocessing

All data were analyzed using the Matlab 2016a open source toolbox Fieldtrip (Oostenveld et al., 2011). Data were segmented 1.5 s before and after the onset of the target word for each trial, including the blank 800 ms period before target word presentation. The segmented data were low-pass filtered at 150 Hz and high-pass filtered at 0.1 Hz and re-referenced to the average of the left and right mastoid. The 50 Hz line-noise component was removed using a discrete Fourier transform (DFT) filter. Remaining strong line noise and muscle artifacts were identified first by visual inspection of amplitude variance over trials and the corresponding trials were removed. Second, artifacts related to eye-movements were removed by means of an independent component analysis (fastICA) (Hyvärinen & Oja, 2000), followed by back projection. Bad channels were repaired by replacing them with the plain average of the nearest neighbors. Third, the resulting data were again visually inspected on a trial-by-trial basis and trials with remaining artifacts were removed. From this procedure and for both groups, 6% of trials were excluded on average from further analysis.

### Event-Related Potential (ERP) analysis

Event-related potentials were investigated to observe N400 and PNP amplitude modulations after target word onset as a function of Constraint (HC, MC, LC), Congruency (congruent, incongruent) and Predictive Validity (high, low). Per condition, pre-processed epochs were low-pass filtered at 35 Hz. Baseline correction was performed using a time-window of −300 ms to 0 ms relative to target word onset. The N400 component was calculated by averaging amplitudes from 250 ms to 600 ms following target word onset. The PNP was calculated by averaging amplitudes from 600 ms to 1000 ms following target word onset. Cluster-based permutation statistics (Maris & Oostenveld, 2007) were used to identify a cluster of channels that resulted from a significant difference in N400/PNP amplitude between levels of the factor Congruency, irrespective of the factors Predictive Validity and Constraint. For subsequent statistical analyses, the average amplitude over the channels belonging to this cluster was extracted per participant and per condition (Constraint x Congruency x Predictive Validity) separately within the N400 and PNP relevant time-window. All statistical analyses on the extracted data were performed in the R software (Core Team, 2019) by fitting a linear mixed effect model (lmer) to the interaction of the conditions (Constraint (within-subject factor): HC, MC, LC; Congruency (within-subject factor): congruent, incongruent; Predictive-validity (between-subject factor): high, low) with participant as random intercept. The estimates of the model were interpreted using R’s type II anova() function with a default treatment coding for contrast estimation. Correction for multiple comparisons was performed using the Tukey method (Tukey, 1949).

### Time-frequency analysis

Time-frequency analysis was performed on a time-window of −1500 ms to 1500 ms relative to target word onset. Power was estimated for a frequency range of 2 Hz to 40 Hz, using a fixed 500 ms sliding Hanning window in time steps of 50 ms, and frequency steps of 2 Hz. No baseline correction was performed on the time-frequency data. Cluster-based permutation statistics (Maris & Oostenveld, 2007) were used to identify a cluster of channels that resulted from a significant difference in alpha (8-12Hz) power between levels of the factor Constraint, irrespective of the factor Predictive-validity. Cluster-based permutation statistics were also used to evaluate significant differences in power across a broad frequency spectrum (2-30Hz) with respect to the factor Constraint, irrespective of Predictive-validity and averaged over a −540 ms to 0 ms time-window relative to target word onset, with respect to the channel clusters identified previously. The relevant time-window was selected based on a previous report on the effects of sentence context constraints on alpha (8-12Hz) power from (Terporten et al., 2019). For visualization only, the alpha (8-12 Hz) specific power modulation over time was plotted by selecting the average power for these cluster-specific channels respectively within a time-window of −1.0 s to 1.0 s relative to target word onset. For subsequent statistical analyses, the average power over the respective channel cluster was extracted per participant and per condition Constraint (HC, MC, LC) x Predictive-validity (high, low) separately, in a time window of −540 ms to 0 ms relative to target word onset. Statistical analyses were performed in the R software (Core Team, 2019) by fitting a linear mixed effect model (lmer) to the interaction of the conditions (Constraint (within-subject factor): HC, MC, LC; Predictive-validity (between-subject factor): high, low) with participant as random intercept. This was done separately for the data of the frontal and parietal channel selection. The estimates of the model were interpreted using R’s type II anova() function with a default treatment coding for contrast estimation. Correction for multiple comparisons was performed using the Tukey method (Tukey, 1949). *P*-values smaller than 0.05 were considered significant.

## RESULTS

### Behavioral performance

The accuracy of the answered questions confirmed that participants were paying attention to the content of the presented sentences. The overall accuracy measures show a mean performance of 91% (SD = 5.61%), 84% (SD = 6.97%) and 89% (SD = 6.39) for the HC, MC and LC sentences respectively for the high predictive validity group. The means of the low predictive validity group were 83% (SD = 7.32%), 84% (SD = 3.52%) and 91% (SD = 4.01) for the HC, MC and LC sentences respectively. Accuracy did differ as a function of Constraint (F(2, 62) =4.8, *p* = .01) but not as a function of Predictive Validity (F(1, 62) = 2.33, *p* = .13). A significant interaction was found between the factors Constraint and Predictive Validity (F(2, 62) = 4.8, *p* = .01). Post-hoc contrasts reveal that this interaction is driven by a significant difference in accuracy between MC and LC sentences for the low predictive validity group (*p* = .003).

### N400 amplitude modulation after target word onset

In this study we were interested in dissociating the effect of predictive validity and context constraint on the N400 amplitude. The predictive validity was manipulated by altering the percentage of occurrences of (in)congruent target words. This was expected to influence the validity of the sentence context constraints and therefore the degree to which the target word will be predicted. From previous literature, we expected a gradual decrease in N400 amplitude with an increase in sentence context constraints (Kutas & Federmeier, 2011; Terporten et al., 2019). Collapsed over Predictive-validity and Constraint, the N400 amplitude was significantly modulated by Congruency as shown by the cluster-based permutation statistics (p = .003, Fig. 2). The statistics revealed a channel cluster over central-posterior electrode sites (Fig. 2). Consistent with our expectations, the N400 amplitude within the time window of 250 ms to 600 ms after target onset was reduced for congruent target words as compared to incongruent ones (Fig. 3, shown for Predictive-validity). For congruent target words, the N400 amplitude was further gradually reduced with an increase in context constraints (Fig. 3). HC displayed the strongest reduction in N400 amplitude, followed by MC and LC. The gradual decrease with the degree of constraint for congruent target words was not apparent for incongruent target words.

**Figure 2:**
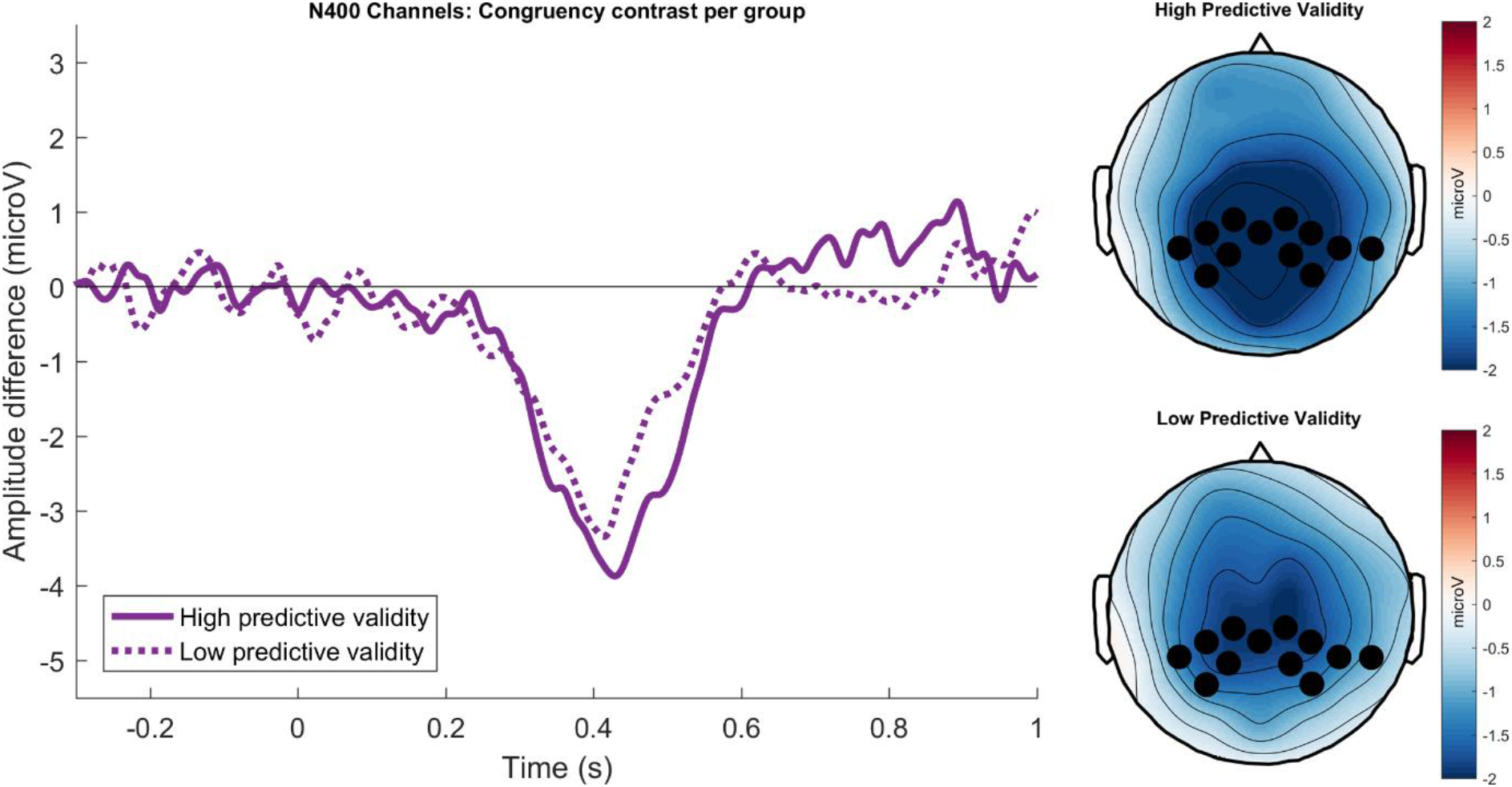
N400 effect comparison between predictive validity groups, averaged over constraints. Left) The N400 effect for the time window 250 ms to 600 ms for each group, as averaged over sentence context constraints. Right) The topography of the N400 effect and the sensor selection based on the top 20% of t-values from the cluster-based permutation statistics.

**Figure 3:**
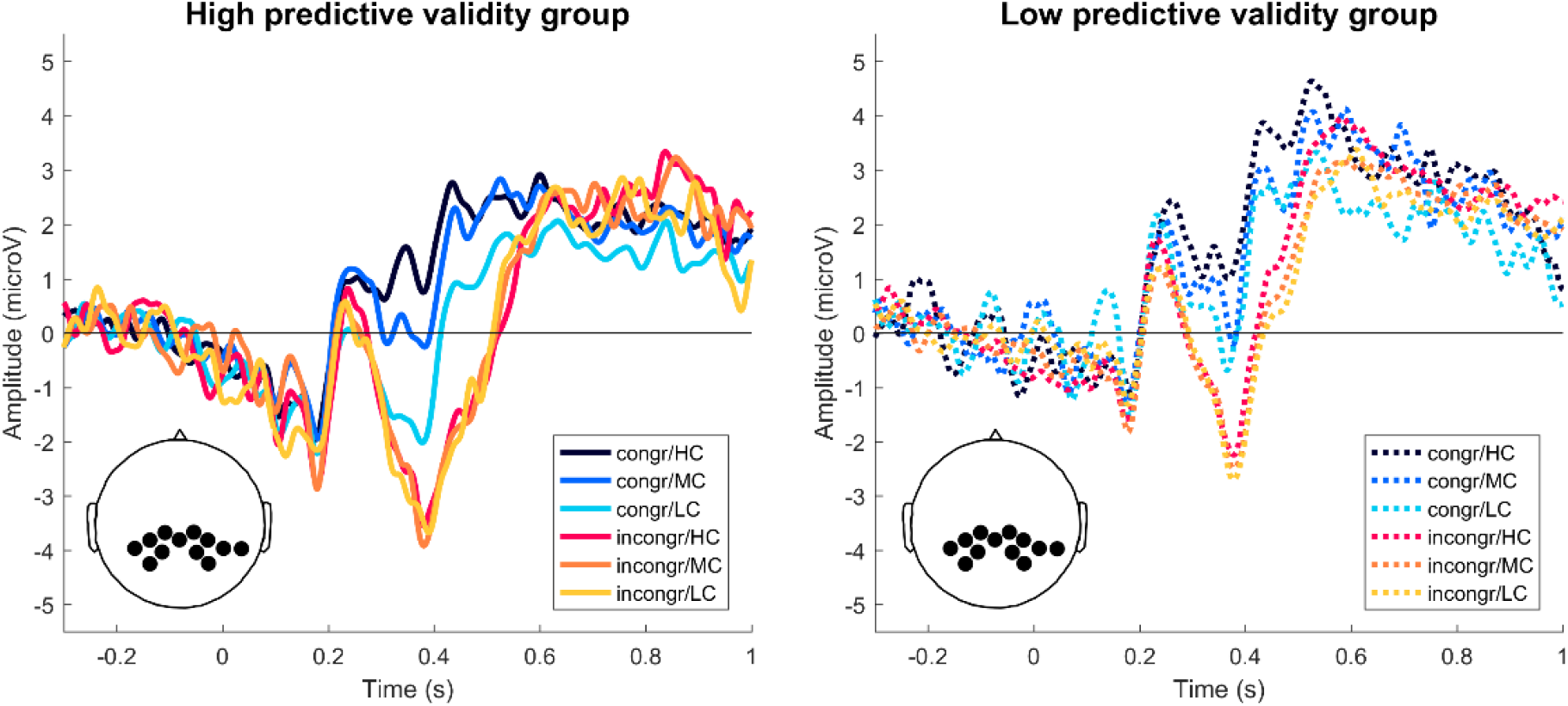
ERP amplitude modulations split by Predictive Validity. Per group, incongruent target words (red colors) resulted in a stronger N400 amplitude than congruent target words (blue colors), within a time-window of 250 ms and 600 ms after target word onset. The effect of context constraints was only observed for congruent target words, for both groups. The groups tend to differ with respect to the overall effects of congruency and constraints onto the N400 amplitude.

The overall patterns of the N400 amplitude modulations as a function of Constraint and Congruency were similar across predictive validity groups (Fig. 3). The ANOVA statistics revealed no significant three-way interaction between Constraint x Congruency x Predictive Validity (F(2, 335) = 0.8, p = .45), no significant interaction between Congruency x Predictive-validity(F(1, 335) = 2.16, p = .14) and no significant interaction between Constraint x Predictive-validity(F(1, 335) = 0.36, p = .7). There is a marginal main effect for the factor Predictive-validity Group (F(1, 67) = 3.83, p = .050). The manipulation of the predictive validity marginally shifted the N400 amplitude across congruency and constraint (Fig. 3). The low predictive validity group displayed overall more positive N400 amplitude modulations as compared to the high predictive validity group (Fig. 3). The ANOVA statistics indicated a significant interaction between the factors Constraint x Congruency (F(2,335) = 4.0, p = .019) and significant main effects of Constraint (F(2, 335) = 8.73, p < .001) and Congruency (F(2, 335) = 120.59, p < .001). Pairwise comparison of the Constraint x Congruency interaction revealed that the effect was driven by a significant difference between LC vs HC (p < .001) and LC vs. MC (p = .011), but not MC vs. HC (p > .100) for congruent target words. The results are in line with the current literature showing that the N400 amplitude is modulated by the constraints of the preceding sentence context when the target word is congruent, not when it is incongruent (Kutas & Federmeier, 2011).

### PNP amplitude modulation after target word onset

We also investigated the effect of predictive validity on the PNP amplitude. Irrespective of Predictive-validity or Constraint, the PNP amplitude was only marginally modulated by Congruency as shown by the cluster-based permutation statistics (p = .076). A channel cluster is revealed over left posterior electrode sites (Fig. S1). Because of a trend in PNP amplitude difference for the factor Congruency, we still performed subsequent statistical tests to explore the data. Overall, the PNP amplitude within the time window of 600 ms to 1000 ms after target onset was reduced for congruent target words as compared to incongruent ones (Fig. S3, shown for each group). For both, congruent and incongruent target words, the PNP amplitude was not gradually modulated by context constraints (Fig. S2). The overall patterns of the PNP amplitude modulations as a function of Constraint and Congruency are similar across predictive validity groups (Fig. S1). The ANOVA statistics revealed no significant three-way interaction between Constraint x Congruency x Predictive-validity (F(2, 335) = 1.31, p = .27), no significant interaction between Congruency x Predictive-validity (F(1, 335) = 0.47, p = .49) and no significant interaction between Constraint x Predictive-validity (F(1, 335) = 0.10, p = .9). The ANOVA statistics indicated no significant interaction between the factors Constraint x Congruency (F(2,335) = 2.4, p = .093) and significant main effects of Congruency (F(2, 335) = 30.77, p < .001), but not Constraint (F(2, 335) = 0.588, p < .56).

### Alpha power modulations before target word onset

Alpha (8-12 Hz) power modulations before target word onset were investigated to study the influence of context constraint and predictive validity on brain states before the occurrence of the target word. We expected alpha power to be modulated by context constraints (Piai et al., 2014; Rommers et al., 2017; Terporten et al., 2019; Wang et al., 2017), but did not expect a linear relationship between context constraint and alpha power (Terporten et al., 2019). Based on Terporten et al. (Terporten et al., 2019) we further expected to observe this effect to be strongest over fronto-parietal electrodes.

Irrespective of Predictive Validity, alpha power is only marginally modulated by Constraint as shown by the cluster-based permutation statistics (p = .085). A channel cluster is revealed over bilateral frontal electrode sites (Fig. 4A). Based on previous results (Terporten et al., 2019), we analyzed the effect of context constraints on neural oscillatory power ranging from 2 Hz to 30 Hz over a time-window of −540 ms to 0 ms prior target word presentation. for a broader frequency spectrum (2-30Hz) and averaged over the earlier indicated channel cluster, revealed a peak in F-values around the alpha frequency band (Fig. 4B). This suggests that alpha oscillations were the most sentively modulated by sentence context constraints as compared to other lower frequency bands. Because of a trend in alpha power differences for the factor Constraint, we still performed subsequent statistical tests to explore the data. Pairwise comparisons revealed that the effect of Constraint was driven by a significant difference between the levels MC vs. HC (p = .008) and LC vs. HC (p = .018), whereas alpha power for the levels LC vs. MC (p = .963) did not significantly differ from each other. The alpha power decrease was found to be stronger for HC, followed by LC and MC. With respect to the data extracted from the cluster, the ANOVA statistics for the time-window −540 ms to 0 ms relative to target word onset indicate no significant interaction between the factors Constraint x Predictive Validity (F(2, 136) = 0.459, p = .633) and no main effect of Predictive Validity (F(1, 68) = 1.322, p = .254). (Fig. 4C).

**Figure 4:**
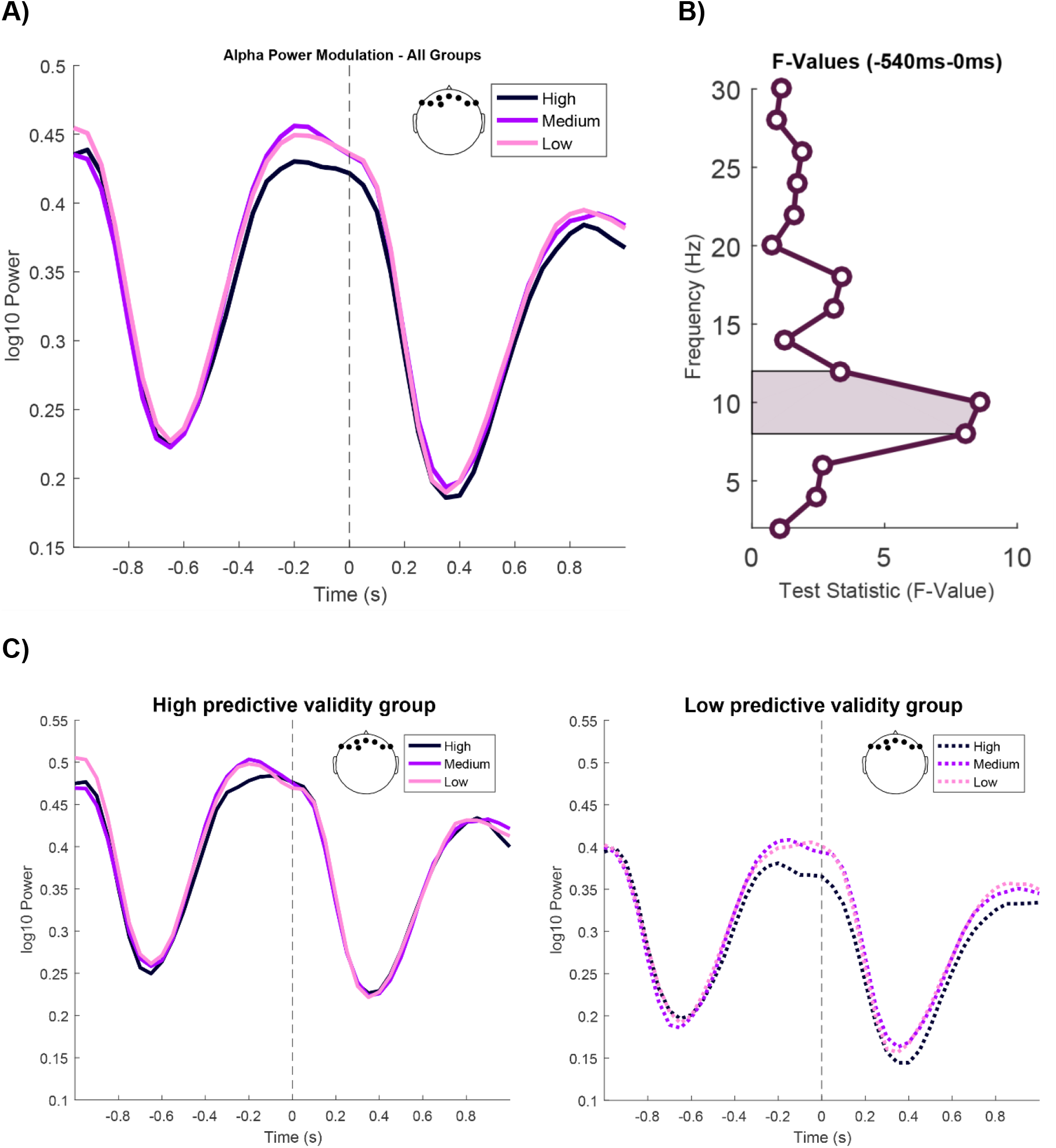
Alpha power modulations as a function of context constraints and Power modulations across a broad frequency spectrum. **A)** Alpha power modulations as a function of context constraints averaged across predictive validity groups. Pre-target word (−540 ms to 0 ms) alpha power is modulated by sentence context constraints. HC contexts induce a stronger alpha power decrease than MC or LC sentence contexts. This effect is most pronounced over frontal electrodes. **B)** Power modulations across a broad frequency spectrum as a function of sentence context constraints, irrespective of Predictive Validity. The shaded part marks the alpha frequency band of interest (8-12Hz). The power spectrum displays a peak in the effect of constraint around the alpha (8-12Hz) frequency band, suggesting that alpha as compared to other frequency bands serves as a cognitive marker that is specific to variations in sentence context constraint. **C)** Alpha power modulations as a function of sentence context constraints, split by Predictive Validity. The pre-target (−540 ms to 0 ms) word alpha power modulations suggest an interaction between Predictive Validity and context constraints, as the effect of constraints appears to be stronger for the incongruent group. This interaction however does not reach significance.

## DISCUSSION

The current study addressed the issue of whether processing of previous semantic context could be affected by the predictive validity of contextual information, for sentences with three degrees of contextual constraints. The validity of these predictions was manipulated group-wise, by changing the proportion of sentence final (target) words that were congruent to the previously established sentence context. Pre-target alpha oscillatory power, and post-target N400 amplitude modulations were investigated as functional markers for the interaction between context constraint and predictive validity.

N400 amplitude was modulated by both the congruency of the target word with its preceding context, as well as the amount of sentential context constraints. However, the N400 did not significantly differ between the predictive-validity groups. For both groups, incongruent target words resulted in a stronger (more negative) N400 amplitude than congruent target words, which is in line with classic N400 findings (Kutas & Federmeier, 2011). A graded difference in N400 amplitude as a function of sentence context constraints was only found for congruent target words, which also confirms our expectations (Kutas & Federmeier, 2011) and replicates earlier investigations of this stimulus material (Terporten et al., 2019). The effects of context constraints and target word congruency on N400 amplitude were not significantly affected by predictive validity.

The robustness of the observed N400 effects across groups speaks against a top-down modulation of linguistic processing as a function of predictive validity and stands in contrast to previous evidence (Brothers et al., 2017, 2019; Lau et al., 2012). Brothers et al. (Brothers et al., 2019) showed an effect of predictive validity on the processing of final words only for highly-constrained sentential contexts. In contrast, we introduced a finer granularity of context constraints, and orthogonalized the effect of linguistic prediction and predictive validity. Additionally, differences in experimental design could have led to differences in how explicit the manipulation of predictive validity is to the participants. In Brothers et al. (Brothers et al., 2019) variations in predictive validity might have been more explicit than in our current approach, instructing participants to either predict or not predict at the start of the experiment. This could potentially mean that if not otherwise explicitly instructed, participants automatically tend to rely on the language input itself rather than on the statistics of its predictive validity. This in turn could have influenced the strategic approach participants applied to achieve the task’s goal. The current results are additionally inconsistent with findings from previous semantic priming paradigms (Lau et al., 2012), which demonstrated that the proportion of valid predictions modulated the N400 amplitude for highly predictable word pairs. We speculate that the process underlying linguistic predictions created in semantic priming paradigms is inherently different from the underlying process required for the current experiment. While semantic priming may rely on associations (Megan A. Boudewyn et al., 2012; Brothers et al., 2017; Kuperberg et al., 2010; Lau et al., 2012), the predictions generated in the current experiment would originate from the combinatorial operation of individual word meanings in a sentence.

Next to effects of semantic congruency, context constraints and predictive validity on the N400 time-window, we also explored their potential effects on a later time-window. Post-N400 positivities (PNP) have also been shown to be sensitive for contextual constraints (Brothers et al., 2017; Delong et al., 2011; Federmeier et al., 2007; Van Petten & Luka, 2012). The PNP could be linked to re-analysis of problematic semantic input that relates to the previous context (Kolk et al., 2003; Kuperberg, 2007; Van Petten & Luka, 2012). In a literature overview, van Petten and Luka (2012) note that the PNP can be influenced by semantic congruency and constraint. Their influence however is expressed by different topographies, with semantic (in)congruencies affecting the PNP over parietal electrode sites, and different predictabilities predominantly affecting PNPs over frontal electrode sites. Indeed, increased frontal positivities were found for unexpected but plausible continuations of strongly constraining contexts (Federmeier et al., 2007; Thornhill & Van Petten, 2012). De Long, Quante and Kutas (DeLong et al., 2014), systematically manipulated context constraints and plausibility during sentence comprehension and confirmed the anterior-posterior dichotomy of the PNP. Anomalous as compared to plausible sentence continuations affected posterior PNPs, while plausible but unexpected words affected anterior PNPs as compared to plausible and expected words. The dissociable PNP topographies suggest different underlying neural networks supporting the re-evaluations of linguistic input based on semantic plausibility and constraint. We did not find statistical evidence for an overall effect of congruency on the PNP, irrespective of context constraints and predictive validity. Yet, we were able to identify a trend in the data that is consistent with the literature cited above: stronger positive amplitudes for incongruent than for congruent sentence endings over left lateralized posterior electrodes. For these posterior electrodes we were unable to observe effects of constraint and predictive validity. Based on the literature, posterior effects of congruency on the PNP can be expected, while this is not the case for effects of constraint. However, in our approach we not only manipulated semantic congruency and sentence context constraints, but we also altered predictive validity. It is possible that the interplay between these three factors introduced spatiotemporal overlap of latent components that makes them indissociable by the current study design (Brouwer & Crocker, 2017).

Before target word onset, alpha power was modulated as a function of context constraints, but not as a function of predictive validity. The main effect of sentential context constraints was only found for frontal electrode sites, for the time window −540 ms to 0 ms relative to target word onset as pre-defined from Terporten et al. (Terporten et al., 2019) (Fig. 4A, 4C). The post-hoc contrast for the levels of context constraints revealed stronger power decreases for HC than for MC or LC. While the stronger power decrease for high constraints compared to lower constraints is in line with earlier findings (Piai et al., 2017, 2017; Wang et al., 2017), the current results only partially replicate our previous work that used a fine-grained constraint modulation (Terporten et al., 2019). Based on Terporten et al. (Terporten et al., 2019), we expected the alpha power decrease to be strongest for MC, followed by the other conditions. In line with our previous findings, alpha power was non-monotonically related to context constraints (power decreases for LC and MC did not differ significantly), but this time alpha desynchronization was strongest for HC instead for MC. This again speaks against a direct or linear relation between pre-target alpha activity and target word predictability in line with Terporten and colleagues (Terporten et al., 2019). If the modulations in pre-target alpha power reflected processes underlying linguistic prediction, we would have further expected that alpha power would be modulated by the predictive validity of the context. This is not what we observed: alpha power remained unaffected by the predictive validity of sentences. . These results speak again against the role of alpha oscillations in linguistic predictions, and rather suggest that they could be linked to the engagement of cognitive processes from other domains, including cognitive control (Fedorenko, 2014), working memory (M. C. M. Bastiaansen et al., 2002; Piai & Roelofs, 2013; Sauseng et al., 2005) or attention (Boudewyn & Carter, 2018; Keitel et al., 2019; Kristensen et al., 2013).

The investigation of alpha oscillations was based on Terporten et al. (Terporten et al., 2019). We also explore the effect of context constraint on a wider frequency range, including e.g. the beta (16-20 Hz) and theta (4-8 Hz) frequency bands, as beta (Lam et al., 2016; Lewis & Bastiaansen, 2015; Wang et al., 2017) as well as the theta (Molinaro et al., 2013; Rommers et al., 2017) frequency bands have been linked to linguistic prediction before. Yet, as in Terporten et al, (2019) effects of context constraints were most prominently observed in the alpha frequency band.

In conclusion, the current study investigated whether an interaction between sentential context constraints and linguistic predictive validity has an influence on pre-target alpha power or N400 amplitude. The results indicate that both N400 after target word occurrence, and pre-stimulus alpha power are sensitive to semantic context constraints. However, alterations of predictive validity did not result in a difference of alpha power or N400 amplitude. These results therefore suggest that both N400 and alpha power are not unequivocally linked to the predictability of a target word based on larger contextual information, supporting the view that prediction mechanisms are not always involved during linguistic processing (Huettig, 2015).

## Supplementary Figures

**Figure S1:**
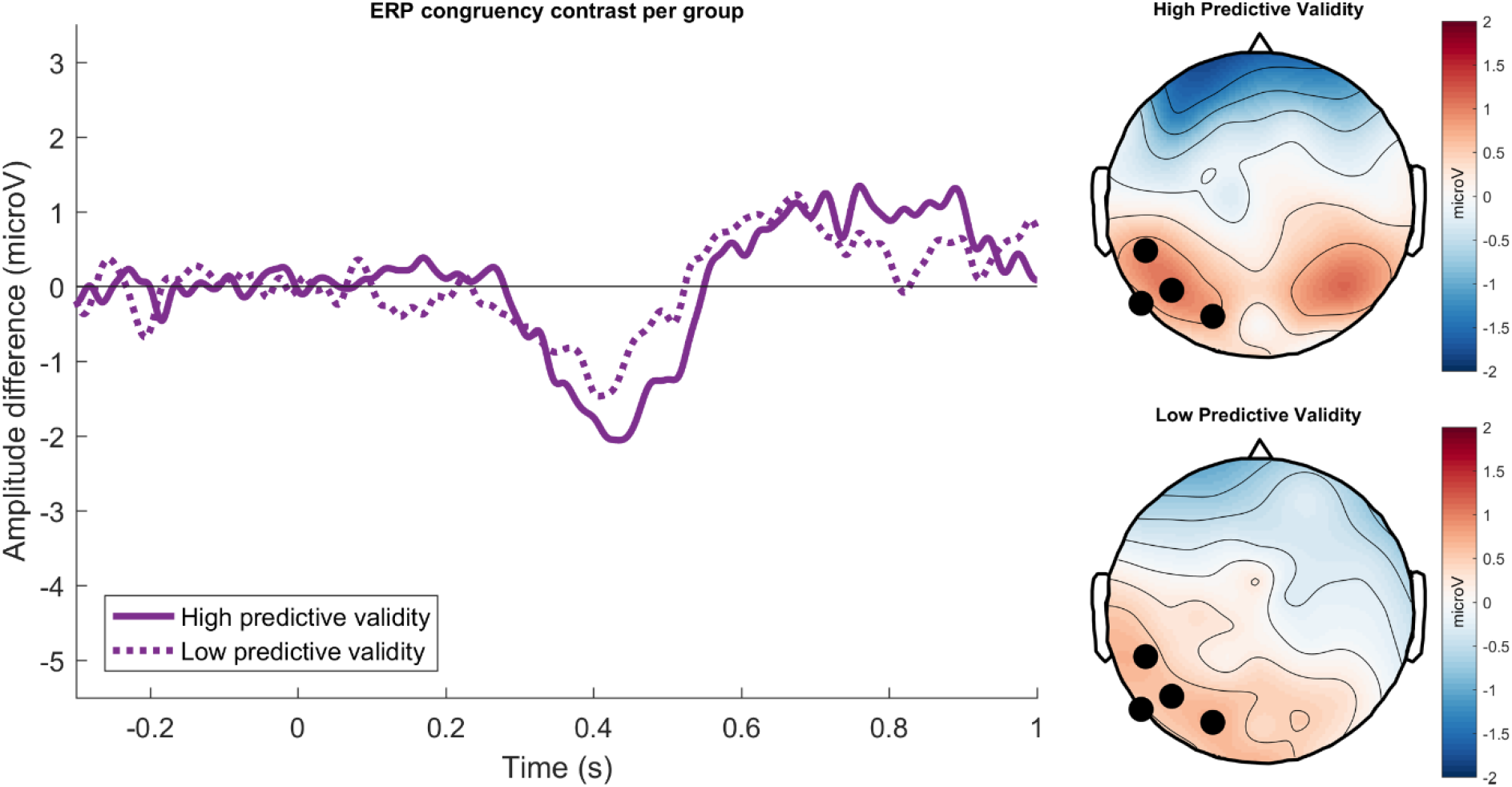
PNP congruency effect comparison between predictive validity groups, averaged over constraints. Left) The PNP congruency effect for the time window 600 ms to 1000 ms as averaged over sentence context constraints. Right) The topography of the PNP congruency effect and the sensor selection based on the corresponding cluster from the cluster-based permutation statistics.

**Figure S2:**
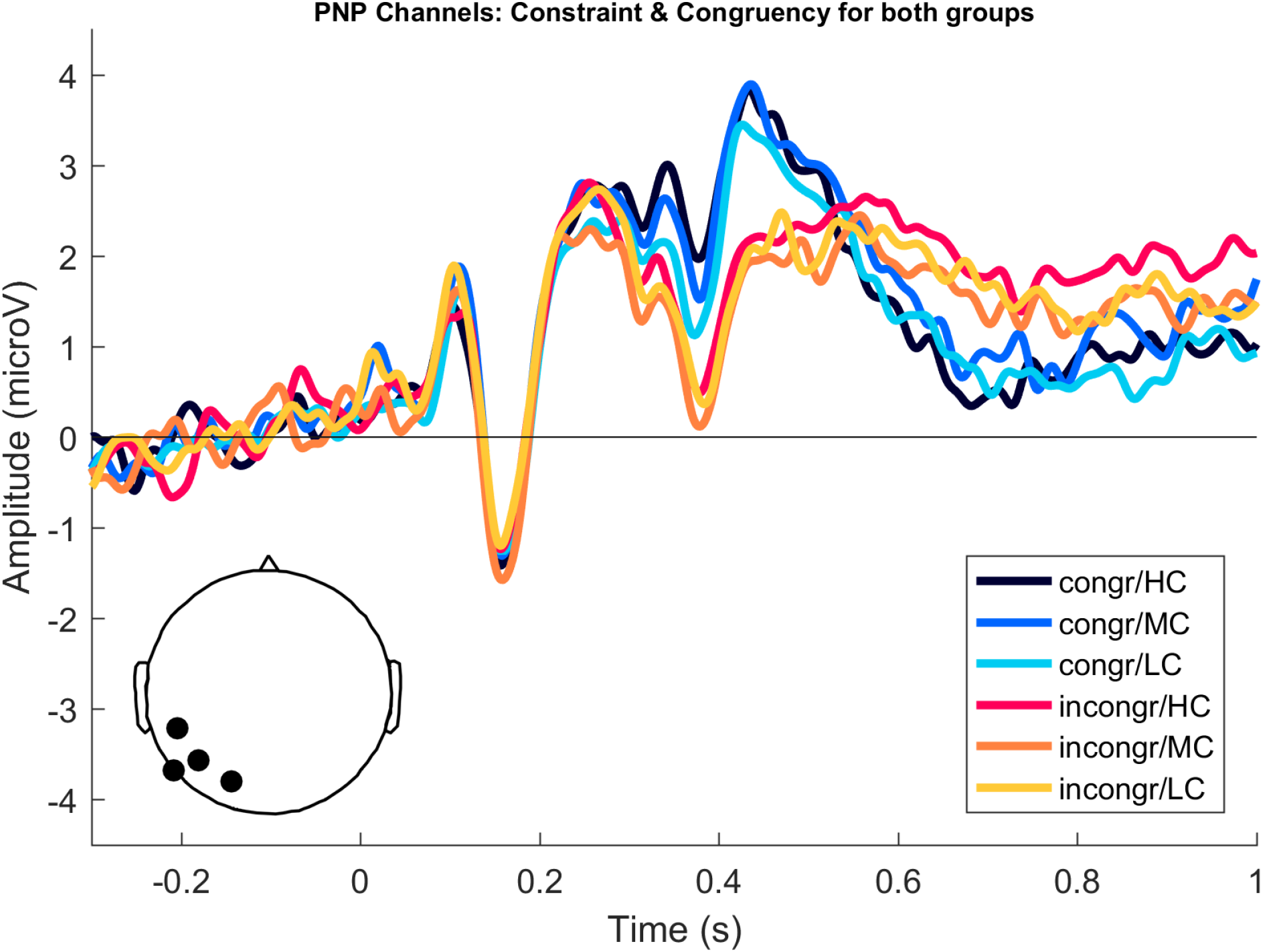
ERP amplitude modulations averaged over predictive validity groups. Incongruent target words (red/yellow colors) resulted in a more positive PNP amplitude than congruent target words (black/blue colors), within a time-window of 600 ms and 1000 ms after target word onset. An effect of context constraints was not observed.

**Figure S3:**
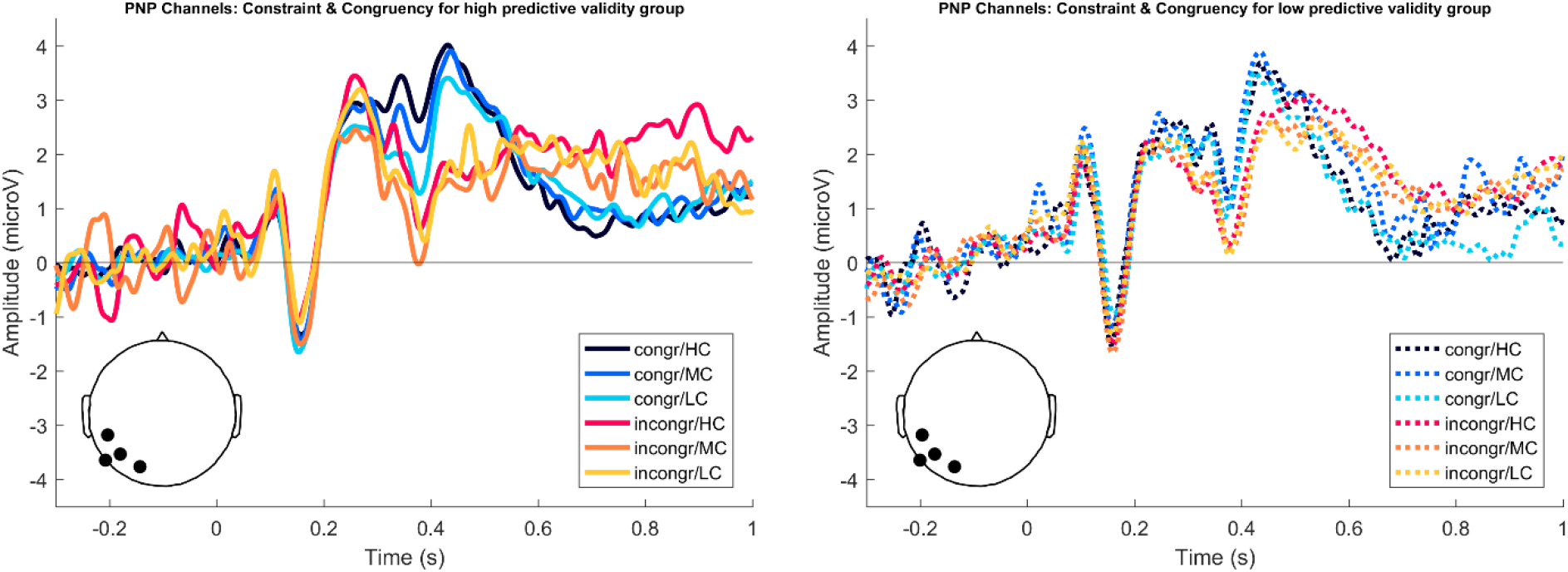
ERP amplitude modulations split by Predictive Validity. Per Predictive Validity group, incongruent target words (red colors) resulted in a more positive PNP amplitude than congruent target words (blue colors), within a time-window of 600 ms and 1000 ms after target word onset. The effect of context constraints was not observed for either group. The groups did not differ with respect to constraint or congruency.

